# Characterization of DNA hydroxymethylation in hypothalamus of elderly mice with postoperative cognitive dysfunction

**DOI:** 10.1101/488007

**Authors:** Jiang Zhong, Wei Xu

## Abstract

Postoperative cognitive dysfunction (POCD) is a common syndrome with perioperative cerebral damage in elderly patients, displaying cognitive impairment and memory loss. Current studies revealed that anesthesia is one of the important causes for POCD occurrence. Recently, Ubiquitin-like with PHD and Ring Finger Domains 2 (Uhrf2) has been reported to play a crucial role in regulating DNA methylation and hydroxymethylation, which are closely connected with memory building and erasure. However, whether narcotic drugs can affect Uhrf2 to impact on DNA methylation and hydroxymethylation in POCD is poorly understood. In this study, we established the elderly POCD mouse model through sevoflurane treatment, and observed the compromised levels of global DNA hydroxymethylated cytosine (5hmC) and Uhrf2 in hippocampus and amygdaloid nucleus compared to non-POCD and control. Furthermore, 5hmC modification on the promoters of the genes associated with neural protection and development, such as GDNF, BDNF, GCR, ACSS2 were reduced in the hippocampus of POCD compared to non-POCD and control groups by MedIP qPCR. Taken together, our findings determined that the loss of 5hmC in hippocampus and amygdaloid nucleus modulated by Uhrf2 suppression might result in the learning and memory ability impairment in POCD.

## Introduction

Postoperative cognitive dysfunction (POCD) is a common syndrome in elderly patients, usually occurred in several weeks or months after operation, representing the dysfunction of central nervous system, such as the cognitive impairment, declining learning and memory ability, information processing disorder and delirium (Pappa et al., 2017). Clinical methodological differences between studies include variable test batteries, lack of control groups, loss of patients during follow-up, and inconsistent intervals between testing periods (Funder et al., 2010). Due to lack of formal diagnostic criteria as well as the subtle cognitive changes, POCD is still difficult to be assessed and diagnosed so far (Monk and Price, 2011).

The causes of POCD in the elderly patients are multifactorial and complicated. One of the potential risk factors responsible for POCD is the use of anesthetic agents. The widely used narcotics currently can be classified into inhalational and intravenous anesthesia. Inhaled general anesthetics such as isoflurane or halothane now have been demonstrated to increase the risk of Alzheimer’s disease (AD) in aging brain (Eckenhoff et al., 2004), and exert neurotoxic effect via caspases-mediated apoptosis (Xie et al., 2008).

One hypothesis proposed is that the epigenetic regulation intervened by anesthetic may be a critical mechanism underlying POCD (Wang et al., 2013). Owing to the similar pathological change course of neurocytes with AD, the general epigenetic alteration on POCD mainly boils down to memory and learning disabilities (Sweatt, 2010). DNA hydroxymethylation is a novel modification based on DNA methylation catalyzed by dioxygenases. The hydroxymethylated cytosine (5hmC) is identified as an intermediate of active demethylation process (Ito et al., 2011; Wu and Zhang, 2017). 5hmC is highly distributed in early embryo, embryonic stem cell (Ito et al., 2010) and nervous system (Kriaucionis and Heintz, 2009; Mellen et al., 2012). The recent studies has revealed that the level of 5hmC reduced 10% after ten-eleven-translocation 1 (Tet1) knockdown can retard the proliferation of neural progenitor cells and impair the abilities of spatial learning and memory (Rudenko et al., 2013; Zhang et al., 2013). Moreover, declining 5hmC modulates transcriptional activity of some genes involved in neurogenesis in AD mice, which also indicates that 5hmC closely connects with memory maintenance (Bernstein et al., 2016; Shu et al., 2016). Besides Tet family, Ubiquitin-like with PHD and ring finger domains 2 (Uhrf2) is currently considered as a novel regulator via its SRA domain (Zhou et al., 2014) to maintain 5hmC level and abilities of learning and memory in brain (Chen et al., 2017b). Here, we hypothesize that the aberrant distribution of 5hmC may be one of the possible molecular causes for POCD occurrence. And which major enzymes contributed to 5hmC metabolism affected by anesthetic in POCD is also poorly understood. Therefore, our study characterize the profiling of DNA hydroxymethylation in central neural system of POCD mice and attempt to reveal the underlying pathogenesis of POCD caused by anesthetic.

## Results

### The overall 5hmC in various nervous tissues display a different extent of change upon POCD

150 mice were treated with 2% sevoflurane for POCD model establishment. Of these mice, 18 were identified as POCD via both Morris water maze test and open field test from two to seven days (Figure 1A, Table 2). Unlike POCD happens most conspicuously on the postoperative 7th day and 3rd month in human (Johnson et al., 2002), we observed that POCD mice started to display the significant loss of memory at 5th day compared to control and non-POCD (Figure 1B, 1C). The mice were sacrificed at 7th day and harvested the brain tissues to detect the global 5hmC via dot blot assay. The presence of differential change of global 5hmC in whole brain lysis before and after POCD was observed (Figure 2A, 2B). Furthermore, hippocampus, amygdaloid nucleus and cerebellum at 7th day were also harvested and observed that the 5hmC was significantly lower in POCD compared to control in hippocampus and amygdaloid nucleus, but no obvious change in cerebellum (Figure 2C, 2D). And we failed to observe any differences of 5mC level between POCD and control (Figure 2E, 2F). Taken together, the loss of 5hmC in hippocampus and amygdaloid nucleus was observed in POCD mice and speculated to be responsible for the cognitive impairment including the loss of abilities of memory, spatial learning and new environmental adaptation.

**Figure 1.**
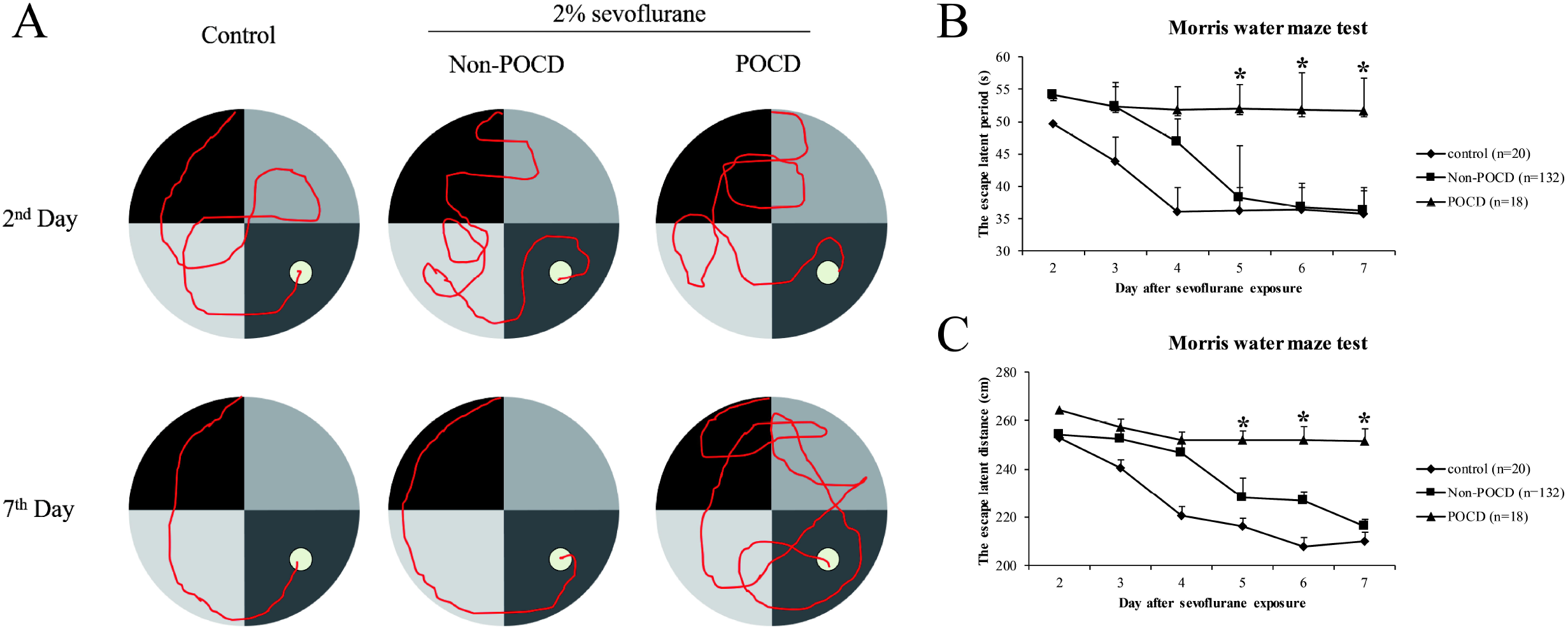
Identification of POCD mice model by Morris water maze test. The movement route (A) and the escape latent period (B) and distance (C) of POCD mice. All data are presented as the mean ± standard error of the mean of 150 mice. *p<0.05 vs. control and non-POCD groups.

**Figure 2.**
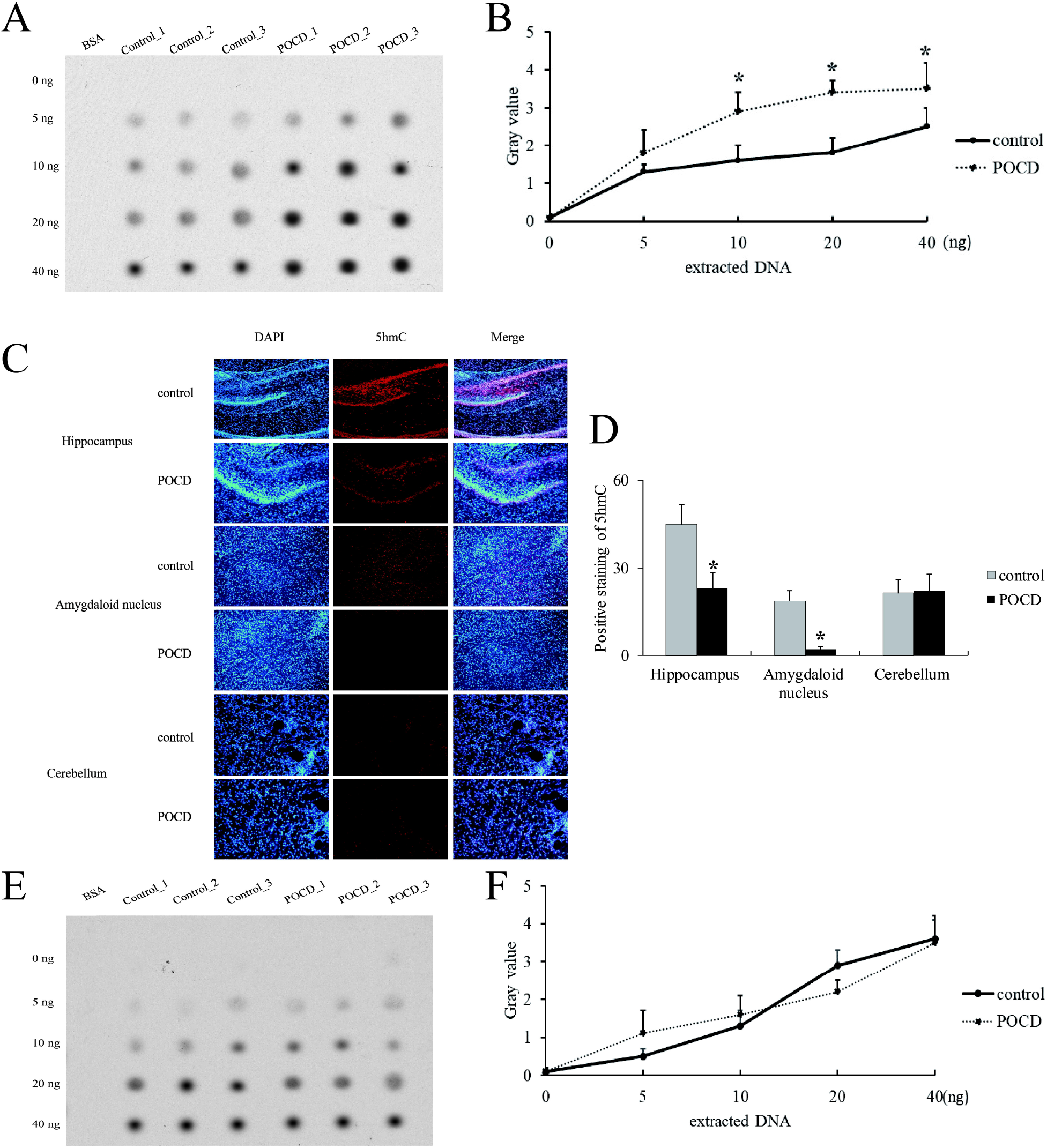
Characterization of 5hmC level in the brain of POCD mice. The global 5hmC level of whole brain using dot blot assay (A) and statistical analysis (B). The 5hmC distribution in hippocampus, amygdaloid nucleus and cerebellum using immunofluorescence (C) and statistical analysis (D). The global 5mC level of whole brain using dot blot assay (E) and statistical analysis (F). All data are presented as the mean ± standard error of the mean of five individual experiments. *p<0.05 vs. control group.

**Table 1.**
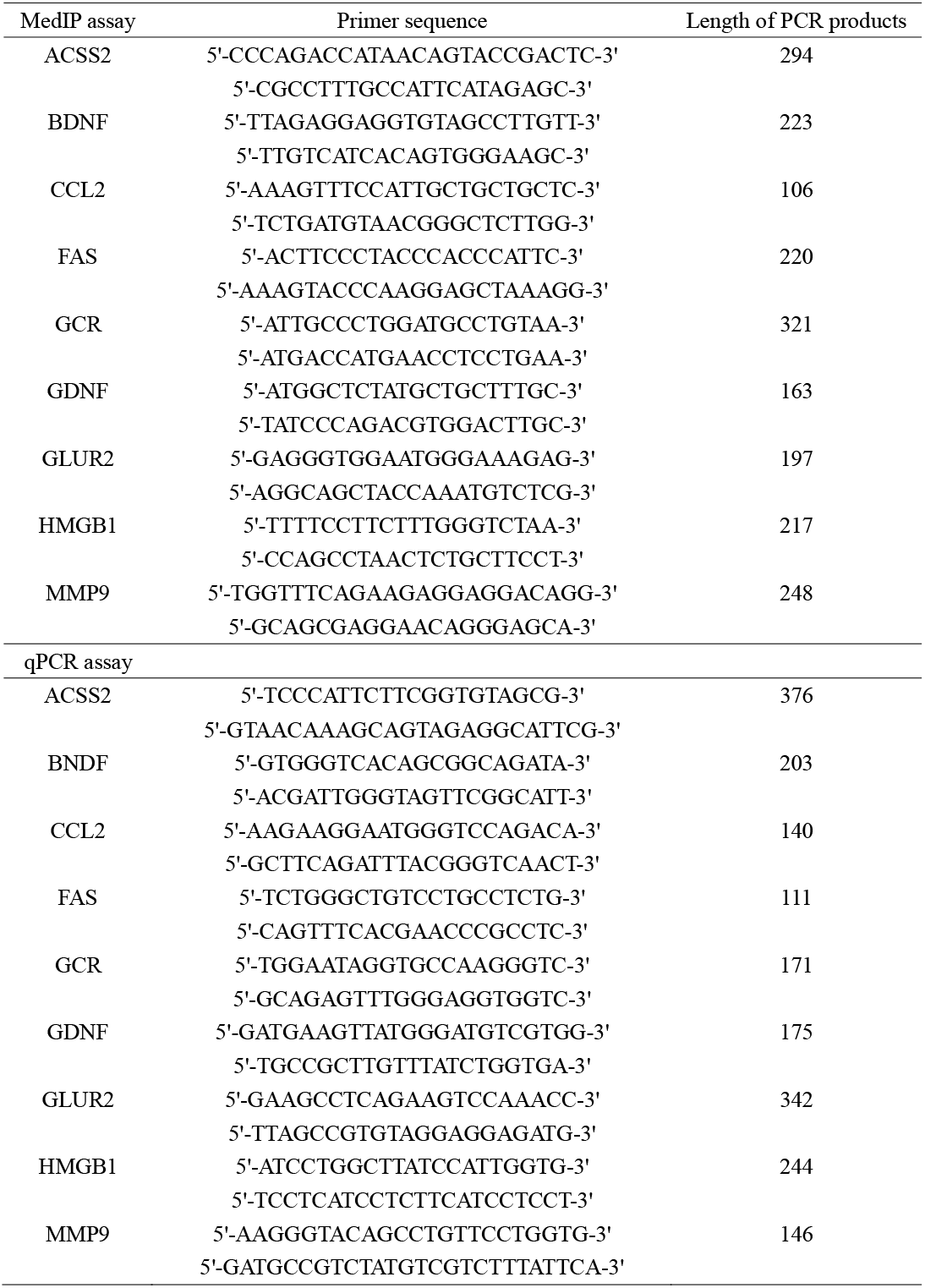
All the primers used in this study.

**Table 2.**
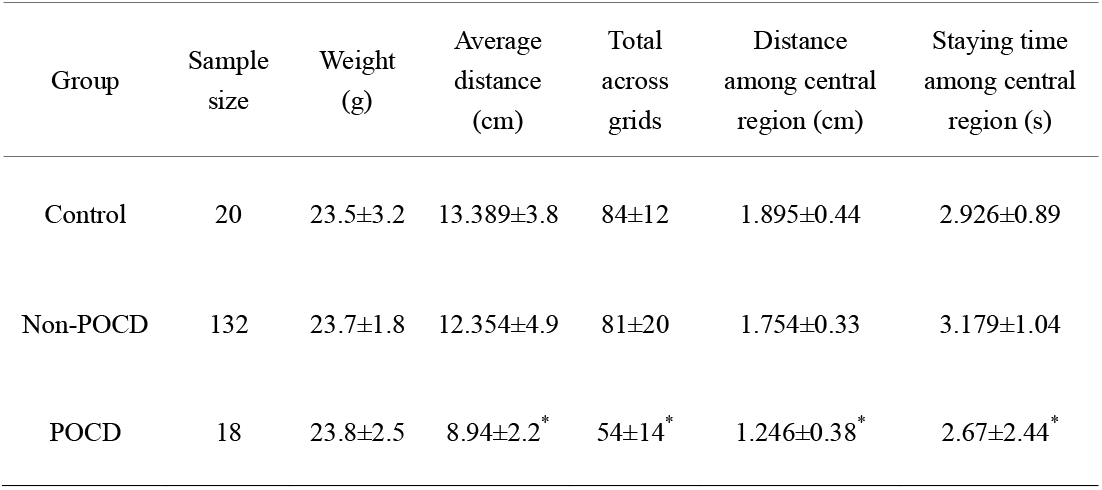
The principle indexes of open field test of POCD mice in this study.

| Group | Sample size | Weight (g) | Average distance (cm) | Total across grids | Distance among central region (cm) | Staying time among central region (s) |
| --- | --- | --- | --- | --- | --- | --- |
| Control | 20 | 23.5±3.2 | 13.389±3.8 | 84±12 | 1.895±0.44 | 2.926±0.89 |
| Non-POCD | 132 | 23.7±1.8 | 12.354±4.9 | 81±20 | 1.754±0.33 | 3.179±1.04 |
| POCD | 18 | 23.8±2.5 | 8.94±2.2<sup>*</sup> | 54±14<sup>*</sup> | 1.246±0.38<sup>*</sup> | 2.67±2.44<sup>*</sup> |

### The loss of Uhrf2 is responsible for 5hmC alteration in hippocampus of POCD mice

To shed some light on the hydroxymethylation impacted by POCD, the enzymes of Uhrf1, Uhrf2, TET1 and TET2 for 5hmC were investigated via western blot. We observed that the protein levels of Uhrf1, TET1 and TET2 exhibited in negligible significance in whole brain between control and POCD, while Uhrf2 displayed a slight down-regulation in POCD (Figure 3A). In consideration about the background noise from the whole brain, Uhrf2 was further investigated in specific regions of brain and observed a suppression in hippocampus and amygdaloid nucleus in POCD compared to control (Figure 3B, 3C), coincident with the change of 5hmC. Collectively, Uhrf2 was suppressed by sevoflurane for hindrance of 5hmC in POCD.

**Figure 3.**
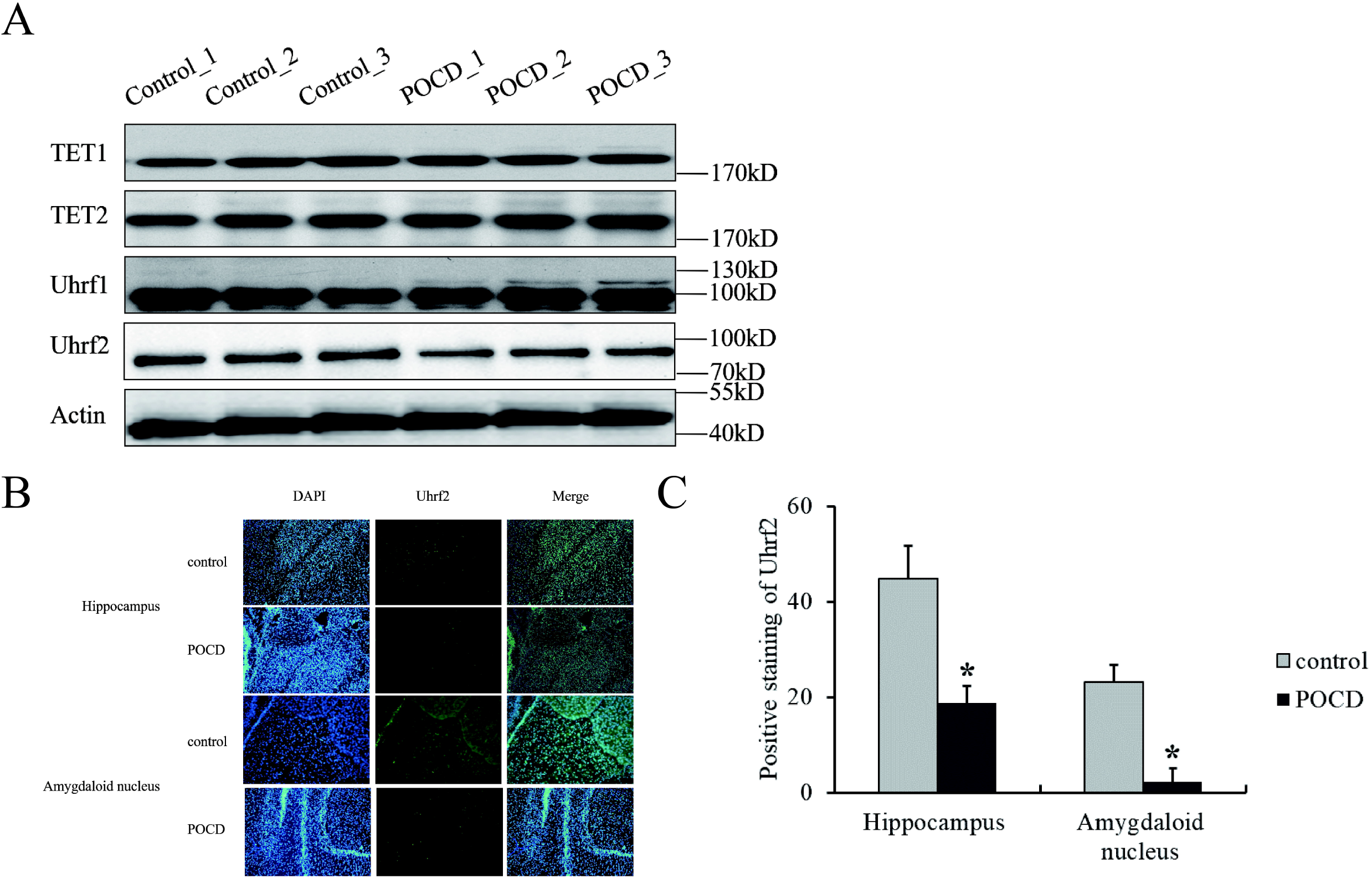
The expression of Uhrf2 in the brain of POCD mice. The enzymes protein levels associated with hydroxymethylation in whole brain using western blot (A). The Uhrf2 protein distribution in hippocampus and amygdaloid nucleus using immunofluorescence (B) and statistical analysis (C). All data are presented as the mean ± standard error of the mean of five individual experiments. *p<0.05 vs. control group.

### POCD results in 5hmC reduction of the certain important genes for neurodevelopment

We supposed that Uhrf2 may be responsible for 5hmC maintenance in hippocampus and amygdaloid nucleus. To further validate the role of 5hmC in POCD, we detected the local 5mC and 5hmC enrichment on the certain genes associated with neurodevelopment through MedIP-qPCR. We observed that GDNF, BDNF, GCR and ACSS2 displayed a reduced 5hmC (Figure 4A) and unvaried 5mC (Figure 4B) level on their promoters in POCD compared to control. Compared with their transcriptional levels (Figure 4C), we observed that 5hmC levels on the promoters of GDNF, BDNF, GCR and ACSS2 could reflect the transcriptional activation of these genes (Figure 4D). Taken together, our results give the evidence that anaesthetics can suppress Uhrf2 to compromise the 5hmC modification on the genes with neuroprotection and proliferation and thereby repress the transcriptional activity.

**Figure 4.**
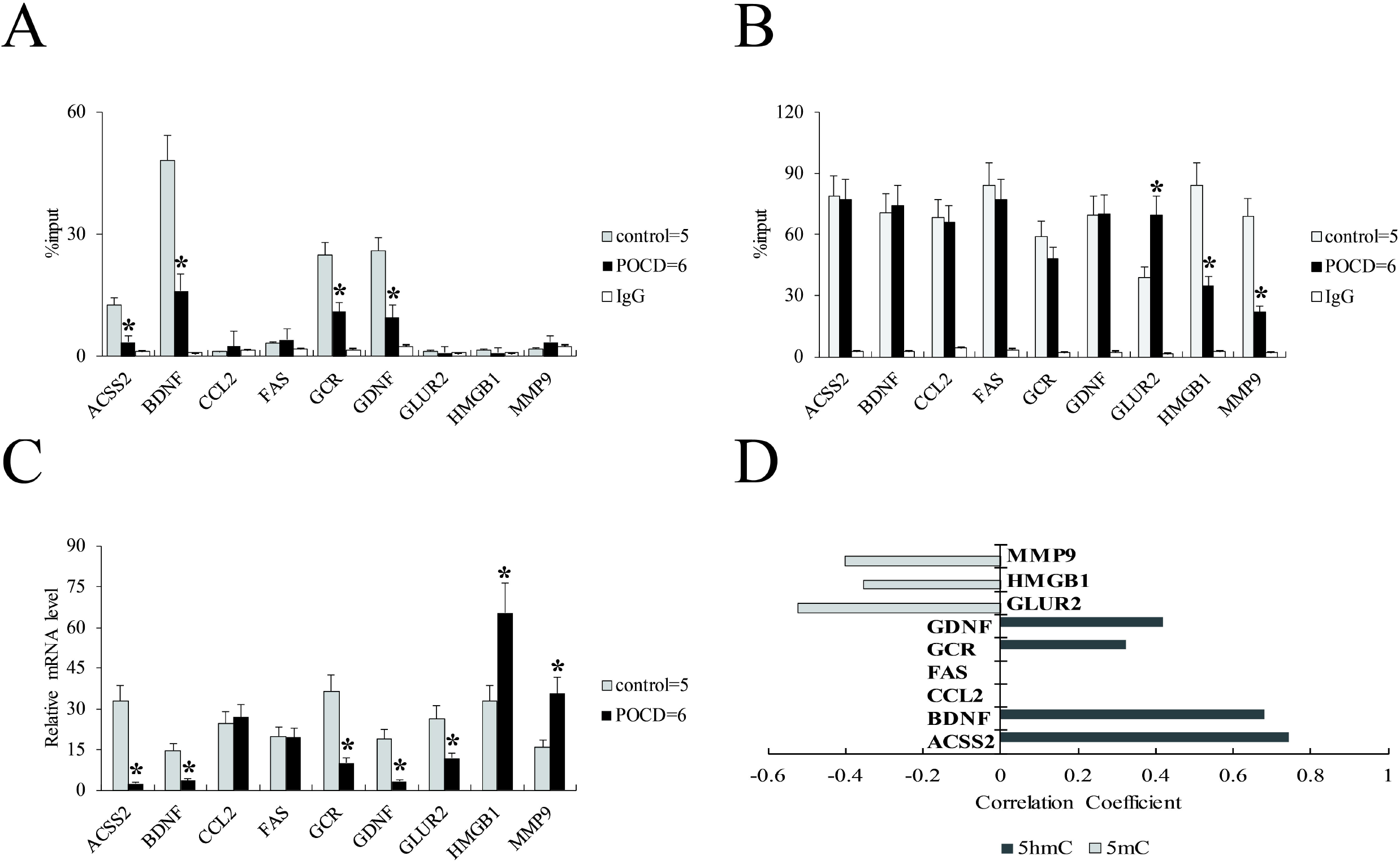
The relationship between 5hmC and gene transcription. 5hmC (A), 5mC (B) enrichment and the transcriptional level (C) in nine candidate genes in control and POCD mice brain. The significant correlation (*p* value less than 0.05) between 5mC or 5hmC enrichment and mRNA level are listed (D). All data are presented as the mean ± standard error of the mean of multiple individual experiments. *p<0.05 vs. control group.

## Discussion

In our study, we used 2% sevoflurane to prepare the POCD model. From the results of ethology, POCD mouse emerged the behavioral and memorial problems in water maze and open field tests. Our observations show that the related data from non-POCD group is similar with normal control (Figure 1, Table 2). Although some phenotypes have been studied to classify early POCD, such as serum proteomics (Zhang et al., 2012), cerebrospinal fluid (Ji et al., 2013), cerebral oxygen saturation (Lin et al., 2013), it is currently still hard to figure out the reasons of individual difference resulting in POCD.

Owing to lack of potent drug for the treatment of POCD in clinical practice, recent study prompts DNA methyltransferase inhibitors which can restore memory-associated transcriptional regulation and improve behavioral memory function in elderly animals (Wang et al., 2013) may be referred to POCD intervention. Nevertheless, there are very few studies investigating the role of epigenetic factors in POCD. Our study bridges 5hmC and POCD, and tries to figure out the role of hydroxymethylation in POCD for the regulation of memory and learning ability maintenance. We observe the global 5hmC level in brain reduced in POCD (Figure 2A, 2B). However, of the different brain regions, 5hmC distributed in hippocampus and amygdaloid nucleus remarkably declines in POCD (Figure 2C), which implies that 5hmC contributes to the memory and learning ability in hippocampus as well as fear emotion control in amygdaloid nucleus. While the presence of monotonous 5hmC level in cerebellum before and after POCD, suggests that the basic associative learning and memory from cerebellum impaired by POCD (Jungwirth et al., 2009) may be independent with accumulation or loss of hydroxymethylation. Consistently, the hydroxymethylation related enzymes such as Uhrf1 and TETs are unchanged (Figure 3A), while the change of Uhrf2 distribution in POCD is consistent with 5hmC both in hippocampus and amygdaloid nucleus (Figure 3B, 3C). The previous study revealed that TET1 knockout mice did not affect overall brain morphology, and furthermore also concluded that TET1 deletion could enhance the consolidation and storage of threat recognition (cued and contextual fear conditioning) and object location memories (Kumar et al., 2015), which is apparently reciprocal with the phenotype of POCD. Therefore, we may rule out the possibility of TET exerting in POCD. Recent study reported that loss of Uhrf2 reduced 5hmC in the brain, including the cortex and hippocampus, but did not change 5mC level, and exhibit a partial impairment in spatial memory acquisition and retention (Chen et al., 2017a), which is strongly confirmed with our results. In a word, our data give the evidence that Uhrf2 is a primary target responding to 5hmC regulation in POCD.

Moreover, we further investigated the relationship between 5hmC enrichment and gene transcription in POCD. In our results, we found two epigenetic ways of the genes transcriptional regulation closely associated with neurodevelopment, to wit, DNA methylation-mediated and loss of DNA hydroxymethylation-mediated gene silencing (Figure 4D). Some genes such as ACSS2, BDNF, GCR and GDNF directly reduce their 5hmC enrichment without any 5mC alteration (Figure 4A), while other genes such as GluR2, HMGB1 and MMP9 present the significant difference of transcriptional levels through changing DNA methylation patterns in promoters (Figure 4B). Herein, we speculate that DNA hydroxymethylation is not the general way to regulate the gene transcription but has some extra condition, such as histones modification, nucleosome package and chromosome architecture.

Overall, we determine that anesthetic lead to the suppression of Uhrf2 and induce the loss of global 5hmC in hippocampus and amygdaloid nucleus, thereby impair the learning and memory ability in POCD. We present 5hmC, which is the important biomarker of memory, and connect with POCD. Our study provides the evidence that Uhrf2 contributed to 5hmC regulation is suppressed by sevoflurane. For POCD evaluation and precaution in perioperation, our study may provide the new index and help develop the new anesthetic in clinic.

## Materials and Methods

### Animal study

All the procedures were followed by the Institutional Animal Care and Use Committee of Fudan University, Shanghai (animal protocol number 2017-32-166). Total 170 40-week-old outbred female C57BL/6 mice purchased from Shanghai SLAC Laboratory Animal Co., Ltd were used in this study. Animals were fed with food and water *ad libitum* freely. 150 mice randomly selected were given 2% sevoflurane for 2h in anesthesia chamber with size of 25cm*13cm*13cm. The other 20 mice were treated normal air as the negative control. Morris water maze test (postoperative 2-7 days) and open field test (postoperative 7th day) were performed to identify the POCD model as indicated (Hovens et al., 2014; Rosczyk et al., 2008). The mice were sacrificed via cervical dislocation and fixed through heart perfusion with 1% paraformaldehyde, and harvested the brain tissues, including hippocampus, amygdaloid nucleus and cerebellum for next studies.

### Dot blot assay

Genomic DNA extracted from tissues (Qiagen, Germany) were dropped 2 μl each on nitrocellulose membrane with 2-fold serially dilution (0, 5, 10, 20 and 40 ng) for dot blot assay. The spots were dried at room temperature and incubated by TBST with 5hmC or 5mC antibodies (1:500 dilution; 1 ng/ml) (Abcam, UK) in 10 ml of TBST for 4 h overnight with gentle shaking. The membranes were washed by TBST three times for 10 min each time in room temperature, followed by rabbit anti-mouse IgG-HRP (1:10000 dilution; 20 ng/ml, Beyotime Biotechnology, China) in 10 ml of TBST for 1 h at room temperature with gentle shaking, and washed again for three times. After that membranes were incubated with 3 ml of ECL Western Blotting Substrate (Beyotime Biotechnology, China) for 5 min in darkness at room temperature and developed the spots.

### Western blot

Brain tissue were homogenized in RIPA buffer solution and then centrifuged at 4 □ at 13,000 rpm for 10 min. The protein quantity in the supernatant was determined using a BCA protein assay kit (Well-bio, China). Equal amounts of protein samples were separated by sodium dodecyl sulfate-polyacrylamide gel electrophoresis (SDS–PAGE) and transferred to polyvinylidene fluoride membranes. The membranes were then blocked by 5% non-fat milk TBS for 90 min and then incubated with the respective primary antibodies of Tet1, Tet2, Uhrf1, Uhrf2 (1:2000, Abcam, UK) and Actin (1:5000, Beyotime Biotechnology, China) overnight at 4□. Membranes were washed in TBST and incubated with rabbit anti-mouse and goat anti-rabbit IgG-HRP (1:10000 dilution; 20 ng/ml, Beyotime Biotechnology, China) at room temperature (RT) for 1 h. Membranes were then treated with an enhanced chemiluminescence detection kit (Millipore), and the intensity of each band was quantified by densitometry.

### Immunofluorescence assay

In brief, mice were fixed by 4% paraformaldehyde in room temperature, and isolated brain tissues with 5 μm sections and washed with PBS, then permeabilized with 0.1% Triton X-100 and blocked with 5% BSA for 30□ min at 37□. Subsequently, slices were incubated with 5hmC and Uhrf2 antibodies at 4□°C overnight. After washing, cells were further incubated with the appropriate Alexa Fluor secondary antibody at 1:20000 dilutions for 30□min at room temperature. After washing, cells were mounted in mounting media with DAPI (Vector Laboratories, Burlingame, CA). The positive staining of 5hmC and Uhrf2 were digitally captured at 400 × magnification and analyzed using Image J software.

### Methylated DNA immunoprecipitation (MeDIP) qPCR assay

For MeDIP assay as previously described (Zhou et al., 2017), extracted genomic DNA was sonicated (90 cycles of 30 s on/30 s off with high power) and incubated with 0.5 ug 5hmC or 5mC antibody or IgG overnight to capture the DNA fragment with 5hmC or 5mC, then washed and harvested for detecting the 5hmC or 5mC enrichment at promoter regions of candidate genes, where the primers setting for qPCR were designed to encompass about 200 bp (Table 1). The qPCR reactions were done using the Fast Universal SYBR Green Realtime PCR Master Mix (Roche, Basel, Switzerland) and in triplicates with the following conditions: 95 °C/30 s, 40 cycles of 95 °C/5 s, 60 °C/30 s. Ct value was analyzed to calculate enrichment using delta-delta methods.

### Realtime PCR

Total RNA was extracted by Trizol Reagent (Invitrogen) according to the manufacturer’s instructions. Before performing reverse transcription, RNA was treated with 5U DNase I (Beyotime) on ice for 10 min to remove the bacteria genomic DNA, and purified by isopropanol and 3M sodium acetate, and washed by 75% ice ethanol, then performed reverse transcription using the QuantiTect Reverse Transcription Kit (Qiagen). qPCR conditions were followed as described above.

### Statistical analysis

Data are presented as means ± standard deviations for three independent experiments. The difference of values was analyzed using Student’s t-test. Pearson correlation analysis is used to evaluate the 5mC and 5hmC correlated with mRNA level. The *p* value less than 0.05 were considered as statistical significance (*, *p*<0.05; **, *p*<0.01).

## Acknowledgements

Not applicable.

## Authors’ contributions

JZ performed experiments and analyzed the data. WX designed the overall project and drafted the paper.

